# Real-time Detection of Acute Lymphoblastic Leukemia Cells Using Deep Learning

**DOI:** 10.1101/2022.10.22.513362

**Authors:** Emma Chen, Mikhail Y. Shalaginov, Rory Liao, Tingying Helen Zeng

## Abstract

Acute lymphoblastic leukemia (ALL) is one of the most common types of cancer among children. It can rapidly become fatal within weeks, hence early diagnosis is critical. Problematically, the ALL diagnosis mainly involves manual blood smear analysis relying on the expertise of medical professionals, which is error-prone and time-consuming. Thus, it is important to develop artificial intelligence tools that will identify leukemic cells from a microscopic image faster, more accurately, and cheaper. Here, we investigate the capabilities of a traditional convolutional neural network (CNN) and You Only Look Once (YOLO) models for real-time detection of leukemic cells. The YOLOv5s model shows 97.2% accuracy for the task of object detection of ALL cells, with the inference speed allowing 80 image frames to be processed per second. These new findings can provide valuable insight in applying real-time object detection algorithms for improving the efficiency of blood cancer diagnosis.

## I. Introduction

Acute lymphoblastic leukemia (ALL) is one of the most dangerous blood cancers known today. Primarily affecting children under 15 years of age, childhood ALL is one of the most common types of cancer among kids. Caused by the excessive proliferation of blood cells, ALL causes the bone marrow to produce many immature lymphocytes, a type of white blood cell (WBC) that defends against infection and illness [1]. When the bone marrow becomes severely impaired, WBC production becomes so rapid that the cells do not properly mature for normal function. As a condition that can rapidly become deadly within weeks, early diagnosis is critical; leukemic cells infiltrate all major body organs, causing them to malfunction or completely fail.

Problematically, leukemia diagnosis in its early stages has challenged researchers and oncologists for years due to the cancer’s initial mild symptoms such as fever, weakness, and bruising. Furthermore, unlike other cancers, leukemia does not create tumors, making it even more difficult to identify. Current manual approaches to blood smear analysis are time-consuming, expensive, and error-prone, relying on valuable resources such as the expertise of trained medical professionals with specialized knowledge of image evaluation [2]. Beyond diagnosis, leukemia patients must endure hundreds of rounds of blood work for blood cell counts and analysis.

There are currently two primary ways to count WBCs. First is through a manual approach in which a blood smear slide is separated into different regions, allowing for a cluster sample to be performed. This approach entails randomly selecting regions of the slide to analyze for an inference calculation of the total number of leukemia cells, which relies on the assumption that the white cells are uniformly distributed across the smear. Although it is possible to count all cells individually without a gridded approach, such a method would be far more time-consuming and is still susceptible to human counting error. The second approach—known as automatic counting—can be more reliable since the technique removes all red blood cells from the sample and automatically counts WBCs with a cell suspension flowing through electronic detectors. However, this approach cannot determine whether cells are damaged or not [3].

As artificial intelligence has been increasingly applied to medicine in recent years, employing machine learning methods can encourage more efficient, precise, and cheap diagnosis and blood work. Early diagnosis is critical to preventing the cancer from spreading and reducing complications, thus raising the survival rate. Not only does increasing the speed of leukemia diagnosis and treatment have the potential to reduce the suffering of hundreds of thousands, but it can also help optimize the healthcare system, allowing doctors and specialists to treat more patients with better quality. The National Cancer Institute predicts that in 2022, the U.S. alone will have 60,650 diagnoses and 24,000 deaths due to leukemia [4]. Our objective is to identify the ALL cells in a blood smear quickly enough for real-time inference. For instance, given a large image of a group of cells, real-time inference could allow doctors to navigate the image while receiving an immediate response on each new section of the image. Exploring many different neural networks, such as a traditional convolutional neural network (CNN) and You Only Look Once (YOLO) models, we strive to make recommendations to researchers to improve leukemia diagnosis and blood cell counting methods.

## II. Materials and Methods

### A. Dataset

The data for this study comes from a public dataset called ALL-IDB, specifically designed for the evaluation of segmentation and image classification algorithms [5]. The microscopic images of blood samples in the dataset were captured with an optical laboratory microscope coupled with a digital camera. The dataset is split into two sections: first is the ALL-IDB1 dataset, which is designed for the purpose of training classification systems. The dataset also contains each image’s blast cell centroid coordinates, if any, indicating the position of ALL lymphoblasts. The second section is the ALL-IDB2 dataset, a collection of cropped area of interest cells from the ALL-IDB1 dataset designed for testing the classification systems’ performances. Since our study focuses on increasing doctors’ efficiency by classifying whole images and detecting individual ALL cells directly from an original lab image, we did not use the ALL-IDB2 dataset, which only zooms in on single cells.

In Figure 1, white blood cells are stained purple, while red blood cells are pink and more plentiful. For our ALL research, only the WBCs are relevant in determining whether the image is healthy or cancerous since those are the cells that can be immature. As seen in Figure 2, the dark center of each WBC is the nucleus, while the lighter outside is the cell’s cytoplasm. Before any models were trained, we resized all images due to different size ratios and computing resource constraints: we formatted the data to 900×1200 for the CNN and 1024×1024 for all YOLO models.

**Figure 1:**
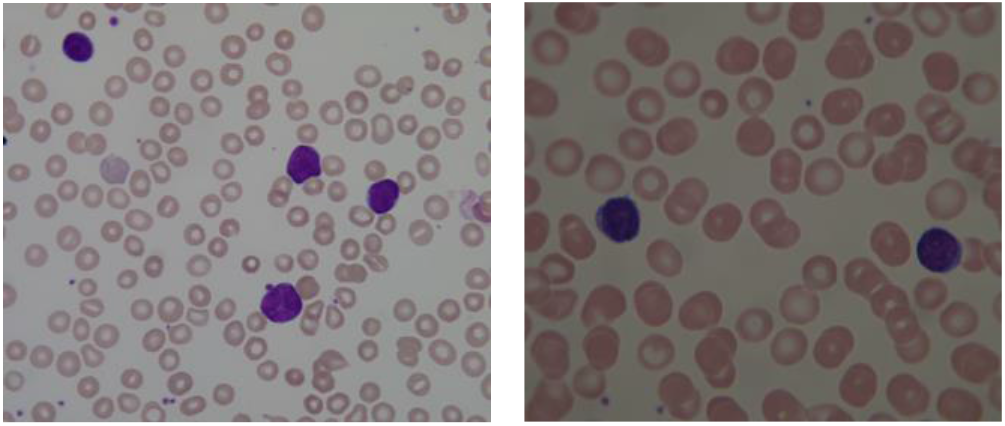
Positive (left) and negative (right) blood smear images from ALL-IDB1. Cancerous and healthy cells can be indistinguishable to the human eye.

**Figure 2:**
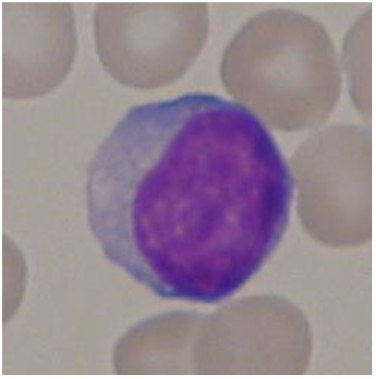
Zoomed in image of a WBC to show detail.

### B. Convolutional Neural Network

The purpose of this study is to identify the leukemic cells in a blood smear image. We started with a classification model aimed at an entire image as healthy or cancerous. After cropping all data images to our target ratio, we separated 15% of the images from the IDB1 dataset for testing and left the remaining images for training. We then built a traditional CNN, which is designed to detect important features and learn adaptively. Since our goal was to reduce the runtime without sacrificing too much accuracy on high-resolution, we created a custom CNN instead of using a pre-trained model.

To compensate for a small dataset, we augmented the data by randomly flipping, rotating, and zooming in the images. After the initial image resizing and rotations, most cancer cells were kept in the image. In order to identify whether a large group of cells contained any ALL cells, we refrained from augmenting the data by zooming too much. Since the images were classified entirely, accuracy values did not change significantly if a few cells were cut out of the image, i.e., all the cancerous-labeled images retained cancer cells so that they could still be classified as positive.

Our CNN model consisted of 3 convolution-pooling layers followed by a dropout layer, a flatten layer, and 3 dense layers with one dropout layer in between. As we crafted the model, we inserted dropout layers between dense layers to prevent overfitting as we deepened the neural network. Training the model was performed within 40 epochs. We stopped adding layers once the training and validation losses stabilized together.

### C. YOLO Models

Once we finished classifying the entire image as positive or negative, we explored object detection models to pinpoint the exact location of leukemic cells with bounding boxes. We employed transfer learning on a publicly available, pre-trained real-time object detection model. The selected models are the YOLO nets, which are known for their incredibly fast speed and advantages in real-time processing. YOLO models have been shown to be one of the best algorithms for object detection, understanding generalized object representation. Our YOLO models were pre-trained on the COCO image dataset, a large-scale dataset of everyday objects and humans. Further, we implemented various YOLO versions for object detection on ALL cells [6].

Using the coordinates from the IDB1 dataset, we annotated all cancer cells for the object detection model. Although the healthy cells were not in a focus of our research, we also annotated all healthy WBCs using our judgment because we had no coordinates. As shown in Figure 3, all leukemic WBCs are enclosed within yellow boxes, while all healthy ones are in pink boxes.

**Figure 3:**
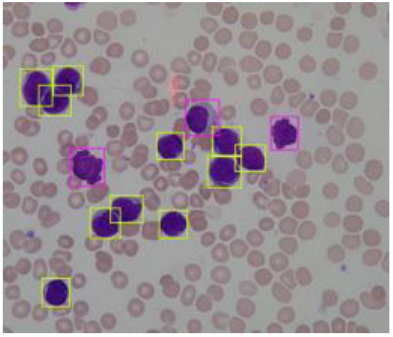
Annotated ground truth blood smear image. Leukemic and healthy WBCs are outlined by yellow and pink frames, respectively.

For the transfer learning, we separated 15% of the images for the test dataset and inputted the remaining training images into the YOLOv5, YOLOv6, and YOLOv7 models, which are different versions of the YOLO model [7]. While YOLOv5 was released in 2020, the YOLOv6 and YOLOv7 were released in June and July of 2022, supposedly outperforming the YOLOv5 in object detection [8, 9]. However, we explored the YOLOv5 more than the YOLOv6 and YOLOv7, given that the version 6 and version 7’s training accuracies were far worse than the YOLOv5 on our ALL dataset—both recent versions continuously identified a cell’s cytoplasm as a completely separate cell. We speculate that the recent YOLO versions’ accuracies were low perhaps because insufficient experimentation, tuning, and revision has yet to be done on them due to their recent publications.

Experimenting with version 5, we researched ALL identification using the YOLOv5 without initial weights and the YOLOv5s and YOLOv5m with pre-trained weights. These models differ in their sizes: the v5s, the small model, has fewer parameters and runs faster than its medium counterpart, the v5m. For instance, the small model contained 213 layers and 7 million parameters, while the medium model had 290 layers and 20 million parameters. As such with fewer parameters, the small model had less weights to adjust during training, which suggests that its accuracy would be weaker than the medium model, but its runtime faster. Finally, we tested the YOLOv5 model, which had no pre-trained weights but the same number of layers and parameters as the small model. Our objective in running this model was to determine whether pre-trained weights from normal, everyday objects would be beneficial in blood cell detection, a potentially very different dataset. Lastly, each YOLO model was trained within 150 epochs.

## III. Results and Discussion

On our test dataset, the CNN classification model performed with a 99.22%. However, it did not identify the location or number of cancer cells like the object detection model, making the CNN much less useful in practice. We focused more on the YOLO models, which could predict bounding boxes and classifications for each WBC cell.

Figures 4, 5, and 6 indicate that the YOLOv5 is capable of reaching a high accuracy during the training process. The model without pre-trained weights performed notably worse than the YOLOv5s and YOLOv5m, likely because the pre-trained weights helped the model identify features and patterns on an abnormal image set as well.

**Figure 4:**
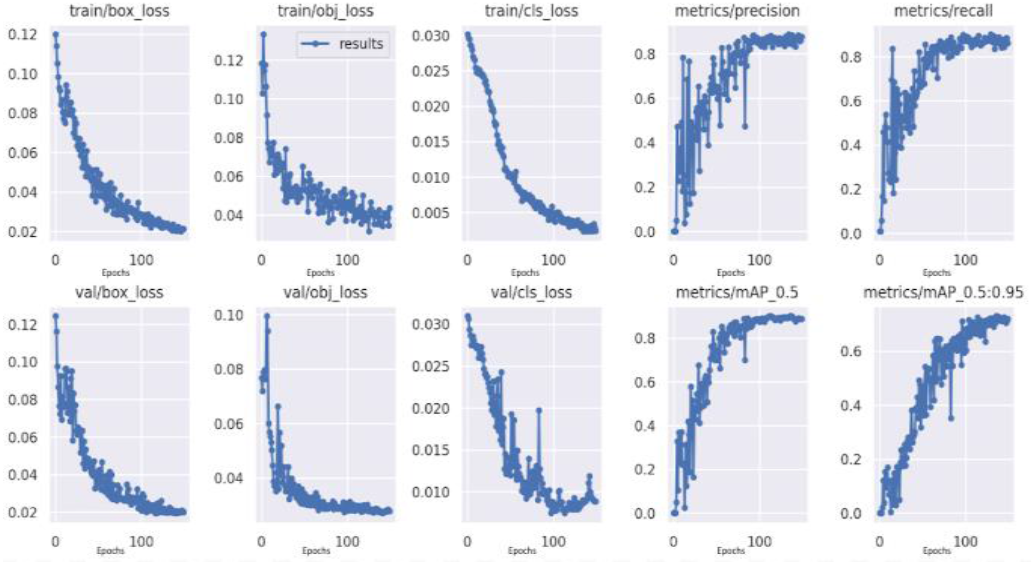
Training graph of YOLOv5s

**Figure 5:**
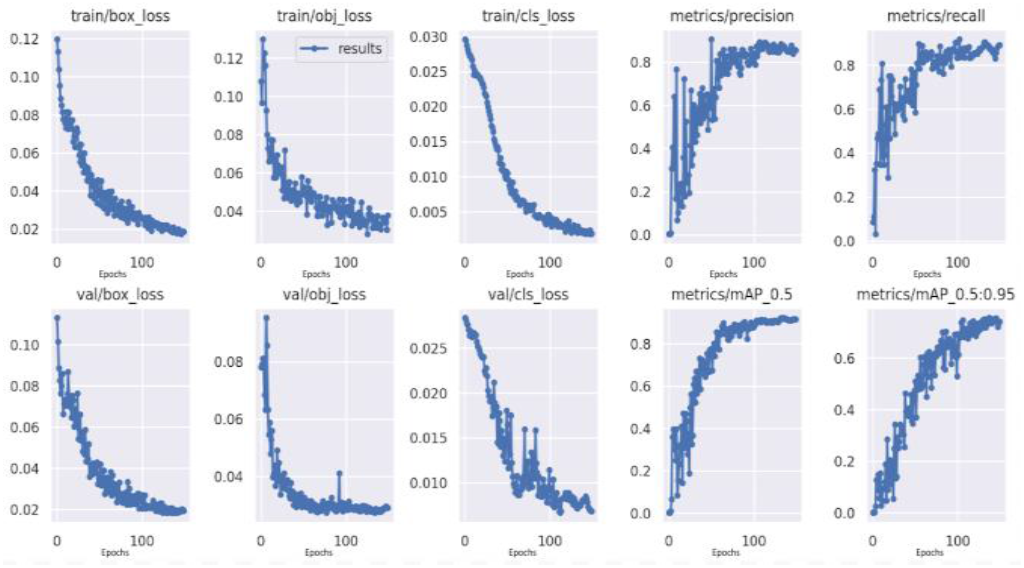
Training graph of YOLOv5m.

**Figure 6:**
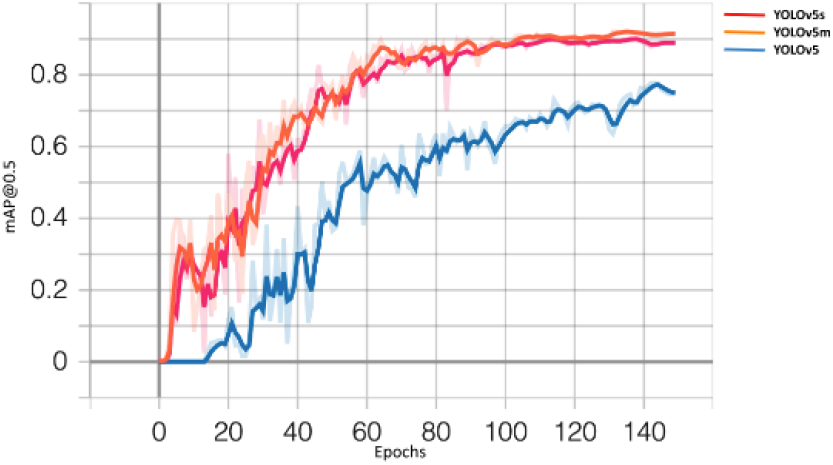
Training graph of all YOLOv5 models depicting that pre-trained weights performed better.

**Figure 7:**
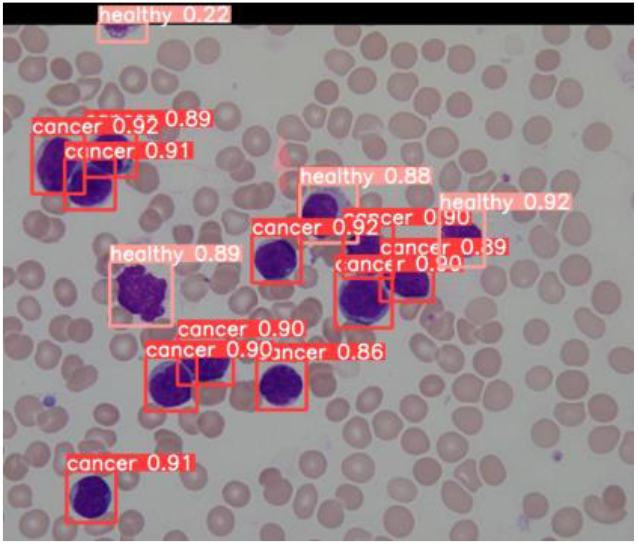
Resulting image from our YOLOv5s’s object detection with confidence levels.

We then evaluated the various YOLO models using 17 images with 126 annotated WBCs to detect. The YOLOv5 models had the highest accuracy and speed.

The metrics in Table 1 are indicators of the models’ performances. Precision (P) measures the quality of a positive prediction made by the model, while recall (R) describes the percent of positives that are classified correctly.

**Table 1:** Performance of YOLO models in detecting ALL cells.

|  | YOLOv5s |  | YOLOv5m |  | YOLOv5 |  | YOLOv6 | YOLOv7 |  |
| --- | --- | --- | --- | --- | --- | --- | --- | --- | --- |
| Cancer(C) / Healthy(H) | C | H | C | H | C | H | C + H (All) | C | H |
| P | .885 | .811 | .936 | .788 | .679 | .736 | N/A | .554 | .425 |
| R | .985 | .787 | .955 | .800 | .970 | .558 | N/A | 1.00 | .814 |
| mAP @0.5 | .972 | .831 | .981 | .853 | .853 | .703 | .644 | .737 | .429 |
| mAP @0.5: 0.95 | .832 | .627 | .861 | .654 | .642 | .401 | .548 | .608 | .330 |
| Inference Speed | 12.3ms |  | 21.9ms |  | 22.0ms |  | 16.0ms | 42.7ms |  |

The mean average precision (mAP) is a metric used to evaluate object detection models. Using a precision-recall curve, the mAP was calculated by the weighted mean of precisions at each intersection-over-union (IoU) threshold. The IoU indicates the overlap of the predicted bounding box compared to the ground truth box. Therefore, a higher IoU indicates that the predicted bounding box is very close to the ground truth box coordinates [10]. The most significant metric is the mAP@0.5 for cancer cells, as we interpret it as the accuracy for the ALL cell location detection. The mAP@0.5:0.95 averages the mAP at each IoU threshold within the interval, but at higher thresholds like 0.95, the margin of error is unnecessarily stringent in practice for our uses; if the predicted bounding box is slightly shifted, a person reading the processed image will likely still be able to interpret which leukemic cell the model is indicating.

We note that the mAP for healthy cells is consistently lower than that of the cancer cells because the healthy cells were self-annotated without precise locations, leaving more room for errors in the ground truth boxes. All the same, our study focuses on identifying the cancer cells in an image, as finding ALL cells can be most useful for oncologists.

As depicted in Table 1, the YOLOv5s and YOLOv5m have significantly higher mAP@0.5 and faster runtimes than the other models. Such speed can be employed for real-time detection: for instance, the YOLOv5s only takes 12.3ms for each inference, meaning that a user could view the model’s ALL identification on 80 unique frames of an image within one second. Finally, our computing machine was the Python 3 Google Compute Engine backend (GPU), for example a Nvidia Tesla P100, from Colab.

## IV. Conclusion

Based on the ALL-IDB dataset of microscopic blood smears, we propose to use the YOLOv5 model for real-time detection of leukemic cells. Specifically, we recommend the YOLOv5s, as the smaller model performs inference with 97.2% mAP accuracy almost twice as fast as YOLOv5m without losing much of accuracy. Our results suggest that object detection models can be more detailed, efficient, and practical than a traditional image classification. As well, the YOLO models could be implemented to report image level classification should such information be warranted. However, given our small dataset, we recognize that our findings should be validated with more field data from hospitals and research facilities. With professional healthy cell annotations, our models could also be modified to assist in blood cell counting in future efforts.

These new findings provide support to continue exploring the possibility of real-time object detection for ALL, which can improve ALL diagnosis and blood work. The YOLOv5 models’ speed and accuracy provide great potential to improve healthcare productivity, and we believe that when implementation is achieved, our results will benefit a great number of people.

## Code Availability

https://github.com/echen053/all-cell-detection

## Acknowledgments

Emma Chen is grateful for the sponsorship and internship training by the Academy for Advanced Research and Development (AARD). This project is partially supported by the Scholarship of Future Scholars of AARD (http://www.ardacademy.org).

